# Disassembly of HIV envelope glycoprotein trimer immunogens is driven by antibodies elicited via immunization

**DOI:** 10.1101/2021.02.16.431310

**Authors:** Hannah L. Turner, Raiees Andrabi, Christopher A. Cottrell, Sara T. Richey, Ge Song, Sean Callaghan, Fabio Anzanello, Tyson J. Moyer, Wuhbet Abraham, Mariane Melo, Murillo Silva, Nicole Scaringi, Eva G. Rakasz, Quentin Sattentau, Darrell J. Irvine, Dennis R. Burton, Andrew B. Ward

**Affiliations:** Department of Integrative Structural and Computational Biology, The Scripps Research Institute, La Jolla, CA 92037, USA; Consortium for HIV/AIDS Vaccine Development (CHAVD), The Scripps Research Institute, La Jolla, CA 92037, USA; Department of Immunology and Microbiology, The Scripps Research Institute, La Jolla, CA 92037, USA; International AIDS Vaccine Initiative - Neutralizing Antibody Center (IAVI-NAC), The Scripps Research Institute, La Jolla, CA 92037, USA; Koch Institute for Integrative Cancer Research, Massachusetts Institute of Technology, Cambridge, MA 02139, USA; Wisconsin National Primate Research Center, University of Wisconsin-Madison, Madison, WI 53715, USA; Department of Biological Engineering, Massachusetts Institute of Technology, Cambridge, MA 02139, USA; Howard Hughes Medical Institute, Chevy Chase, MD 20815, USA; The Sir William Dunn School of Pathology, The University of Oxford, Oxford, OX1 3RE, United Kingdom; Ragon Institute of MGH, MIT and Harvard, Cambridge, MA 02139, USA

**Author notes:** Teaser Statement: HIV Env trimers elicit antibodies that lead to their own destruction.

## Abstract

Rationally designed protein subunit vaccines are being developed for a variety of viruses including influenza, RSV, SARS-CoV-2 and HIV. These vaccines are based on stabilized versions of the primary targets of neutralizing antibodies on the viral surface, namely viral fusion glycoproteins. While these immunogens display the epitopes of potent neutralizing antibodies, they also present epitopes recognized by non or weakly neutralizing (“off-target”) antibodies. Using our recently developed electron microscopy epitope mapping approach, we have uncovered a phenomenon wherein off-target antibodies elicited by HIV trimer subunit vaccines cause the otherwise highly stabilized trimeric proteins to degrade into cognate protomers. Further, we show that these protomers expose an expanded suite of off-target epitopes, normally occluded inside the prefusion conformation of trimer, that subsequently elicit further off-target antibody responses. Our study provides critical insights for further improvement of HIV subunit trimer vaccines for future rounds of the iterative vaccine design process.

## Introduction

HIV infects approximately 1.7 million individuals each year, with an estimated 38 million currently living with the virus (unaids.org). While there are a large number of anti-retroviral therapies available, a vaccine would be the most effective measure for controlling the spread of the virus and best hope for eventual eradication. To that end there are a number of parallel efforts being developed at preclinical and clinical stages that employ a stable, soluble HIV envelope glycoprotein (Env) trimer as a protein subunit immunogen(*1*). In preclinical animal models, variants of these trimers have elicited consistent autologous and sporadic heterologous neutralizing antibody responses against difficult-to-neutralize Tier 2 viruses(*2*–*5*). Although these results are encouraging, there is still much work to be done to elicit broad and potent immune responses. To this end there are significant efforts to understand the types of immune responses elicited by various trimer vaccine candidates in great detail. As such, elucidating on- and off-target (neutralizing and non-neutralizing) antibody responses elicited by these trimers remains a critical component for immunogen redesign and the iterative, rational vaccine design process(*6*, *7*).

Env is an inherently metastable protein because it must undergo drastic conformational changes to drive entry of the virus into the host cell. Broadly neutralizing antibody epitopes are only present on the prefusion conformation of Env, and therefore rational vaccine design is focused on producing this form of Env(*1*, *8*). Most of the current soluble Env trimer immunogens are based on the SOSIP design that includes a disulfide bond which links gp120 to gp41 and an isoleucine to proline mutation in the heptad repeat 1 (HR1) region that stabilizes the pre-fusion conformation of the trimer(*3*, *9*–*11*). Further, truncating the gene at residue 664 improves solubility and monodispersity of the trimer(*12*). Such trimers are designed to antigenically mimic native envelope trimers (Env), including glycosylation(*13*–*16*), while also being soluble and highly stable. Soluble HIV Env trimer immunogen design has increased trimer stabilization to above 75 degrees Celsius(*17*). These highly stable trimers mask internal and undesirable epitopes exposed on gp120 or non-native versions of Env, and exhibit an antigenic profile that preferentially exposes neutralizing epitopes and not non-neutralizing, and otherwise immunodominant, epitopes(*18*).

Despite being highly stable with antigenically optimized surfaces, the HIV Env trimers described above still elicit strong non-neutralizing antibody responses(*6*, *7*, *18*). Hence, the trimers require iterative improvement via testing in animals, characterization of elicited antibodies, mapping of epitopes, and further redesign. To accelerate this process we recently described a new technique, electron microscopy polyclonal epitope mapping (EMPEM), which is a relatively high throughput and comprehensive assay for mapping antibody responses in polyclonal sera(*6*). To conduct EMPEM studies immunoglobulins from the sera or plasma of infected or vaccinated animals are isolated, made into complexes with protein subunit vaccines, and then imaged and reconstructed using single particle electron microscopy(*6*). Because of the speed and information content generated by EMPEM, it has become an increasingly valuable tool for informing immunogen redesign, particularly for identifying on- and off-target antibody responses(*7*).

We previously showed that base binding antibodies appeared after the priming immunization in rabbits and were present, typically at increasing abundance, after subsequent booster immunizations(*6*). Immune responses to soluble Env trimer vaccination have also revealed an abundance of elicited base-binding antibodies(*4*, *7*, *19*). Some estimates for base-directed responses are greater than 90%, which is a highly unproductive skewing of antibodies away from the relevant neutralizing epitopes(*18*). Recently, Moyer et al. attached Env trimers to alum via a C-terminal phosphoserine tag at the base of the trimer in an attempt to mask off-target base antibodies in rabbits(*20*). EMPEM analysis revealed that the Alum-trimer complex did not completely prevent base responses although there was a greater diversity of antibodies bound to neutralizing epitopes outside the base, particularly at the earlier timepoints. These data suggest that blunting of the base response early on increased the diversity of epitopes targeted, including neutralizing epitopes(*20*). Other attempts to mask the base via attachment of Env trimers to nanoparticles have only had limited success(*21*).

While the trimer base is typically an irrelevant and undesirable epitope, a few broadly neutralizing antibodies (bnAb) targeting an epitope near the base have been described. Monoclonal antibody (mAb) 35O22 binds parallel to the viral membrane at the base along a conserved region at the gp120/gp41 interface and neutralizes 62% of 181 pseudoviruses with IC50 <50ug/mL(*22*). Glycans located near the gp120/gp41 interface, including N88, are among the most conserved on Env(*22*). Antibodies such as 3BC315 and 1C2 bind at the gp120/gp41 interface to an overlapping but unique epitope from 35O22. Strikingly, binding of these bnAbs causes the Env trimer to fall apart into its protomer components(*4*, *23*) by disrupting the tryptophan clasp that stabilizes the base of the trimer(*24*). Importantly, all of the known neutralizing base antibodies approach the bottom of the trimer from an angle roughly parallel to the membrane while the non-nAbs approach the base at a steeper angle that would clash with the membrane. This angle of approach makes it relatively easy to differentiate neutralizing from non-neutralizing antibodies via EMPEM.

While trimer degradation is a potential mechanism for viral neutralization, it is confined to a small subset of antibodies described above that meet specific criteria. Here, we demonstrate that there are also classes of base-directed, non-neutralizing antibodies that cause soluble Env trimers to disassemble into protomers. All Envs that we have examined degrade in this manner, and we have observed this phenomenon using sera from both rabbits and NHPs. Most perniciously, secondary antibody responses to internal, non-neutralizing epitopes demonstrate that antibody induced trimer degradation happens in vivo and may further thwart productive immune responses.

## Results

### Highly stabilized Env trimers can degrade into component protomers

Single particle negative stain electron microscopy (nsEM) is a simple and quick technique to assess sample purity and map antibody epitopes on viral glycoproteins(*6*, *25*–*28*). This technique has now become commonplace for evaluating the homogeneity and native-like conformation of HIV envelope glycoprotein (Env) trimers being developed as protein subunit immunogens. In this study, we conducted EMPEM using various serum antibodies elicited by different stabilized Env trimers. Within each set of 2D classes we observed a range of phenotypes including intact trimers bound to fragments antigen binding (Fabs) (Fig. 1) as well as monomeric Env, or protomers, bound to Fabs (Fig. 1B). These protomers are compact and appear to remain in the prefusion conformation, likely due to the engineered stabilizing mutations that include a disulfide bond that links gp120 to gp41 and an isoleucine to proline mutation in the heptad repeat region of gp41 that prevents the conformational transition into an extended helix(*3*, *9*–*11*), as well as others. Given the relative abundance of protomeric species in different Env trimer samples imaged for EMPEM analysis, we undertook a systematic study to further investigate the causes and effects of trimer degradation into protomers.

**Fig 1.**
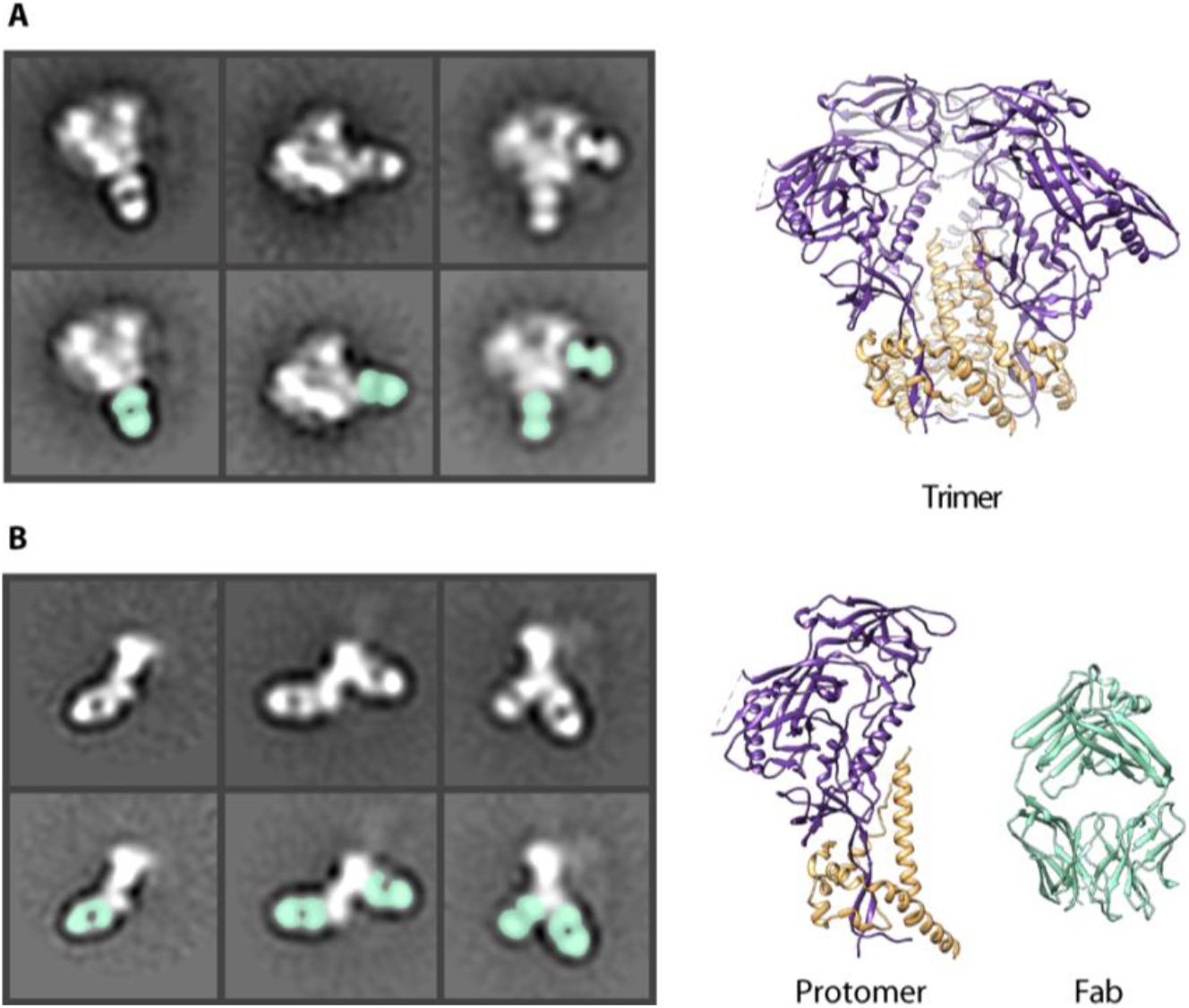
HIV Env protein in negative stain EM. (**A**) Negative stain 2D classes of Env trimer with Fabs bound. Fabs highlighted in seafoam green. CryoEM derived model of Env BG505 SOSIP trimer (6DID). gp120 (purple) and gp41 (orange). (**B**) Negative stain 2D class of Env protomers with Fabs bound. Models of a single Env protomer and Fab (6DID) in seafoam green for comparison.

### Some base-binding antibodies cause trimer degradation

Prior to immunogenicity, antigenicity, or high-resolution structural studies, we routinely evaluate Env trimers by single particle EM to ensure homogeneous distribution of prefusion conformation trimer (Fig. 2A and fig. S1). For epitope mapping, we typically incubate trimers overnight with an excess of Fab (either monoclonal or polyclonal), and the resulting complexes are purified using size exclusion chromatography and deposited onto negative stain EM grids.

**Fig 2.**
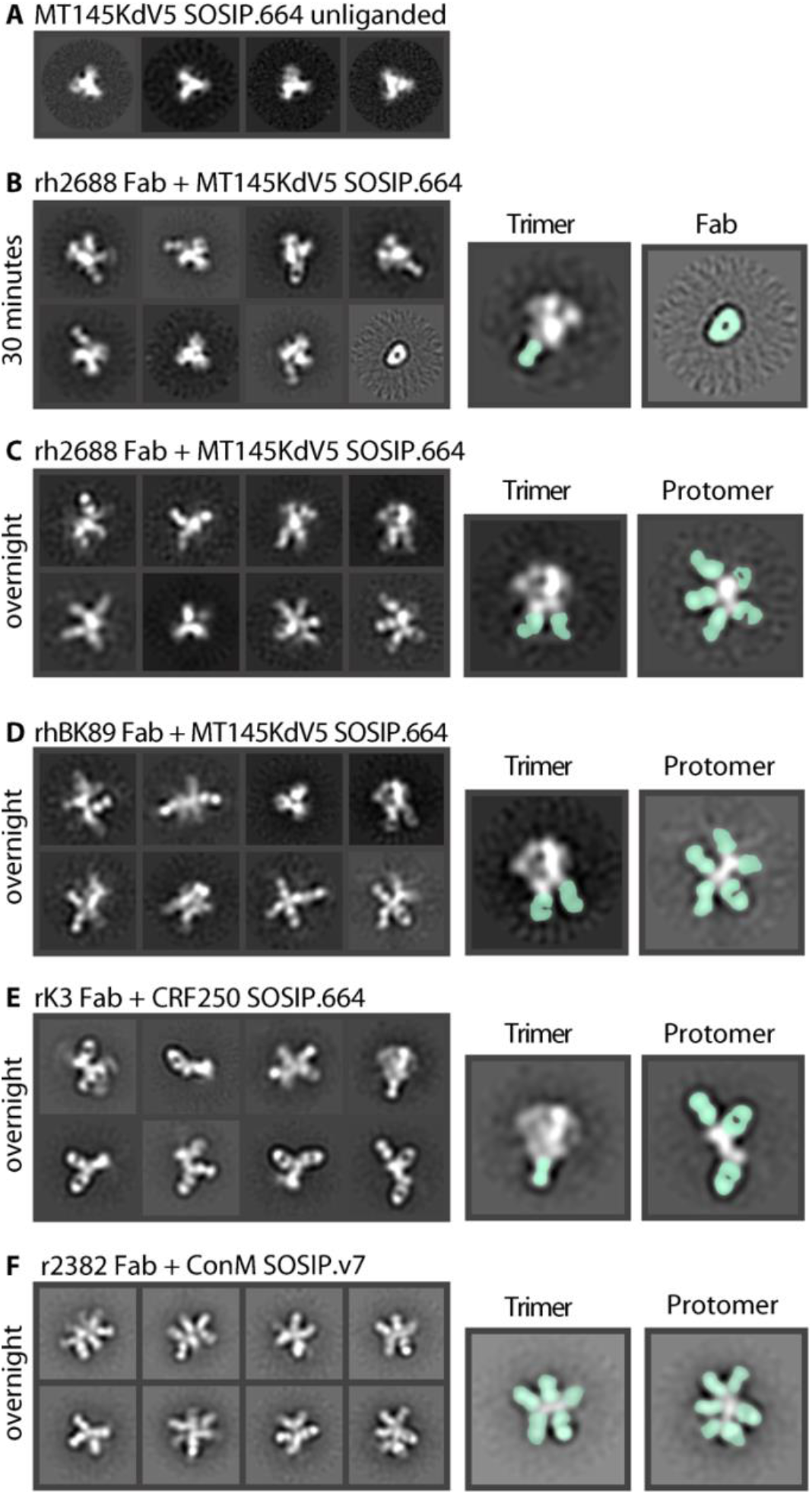
Env degradation after polyclonal Fab incubation. (**A**) Negative stain 2D classes of unliganded MT145KdV5 SOSIP.664 Env trimer. Mostly top views due to lack of tumbling. (**B**) Polyclonal Fab from rhesus macaque 2688 in complex with MT145dV5 SOSIP.664 trimer incubated for 30 mins and placed on nsEM grid. Highlighted classes show intact trimer with Fab bound (seafoam green) (left) and unliganded Fab (right). (**C**) Trimer stays intact within 30 minutes of incubation but will degrade over time allowing secondary Fabs to bind internal neo-epitopes of the protomer. (**D**) Polyclonal Fab from rhesus macaque BK89 in complex with MT145dV5 SOSIP.664 shows a similar outcome as with previous rhesus macaque polyclonal serum. (**E**) Rabbit derived polyclonal Fab in complex with CRF250 SOSIP.664 Env shows trimer degradation and base binding Fabs bound to few intact trimers.(**F**) Rabbit derived polyclonal Fab in complex with ConM SOSIP.v7 Env showing highly decorated Env protomers.

Using Env trimers of various subtypes and serum Fabs from different animal models immunized with soluble Env trimers (rabbits, macaques), we observed a range of trimeric and protomeric Env-Fab complexes (Fig. 2B to F and fig. S2 to S6). Consistent with previous observations, we noted a large percentage of trimers with Fabs bound to the immunodominant base epitope (Fig. 2,B to F). We also observed 2D classes of protomers that were highly decorated with Fabs (Fig. 2,C to F). The extended structure and globular gp120 head of the protomer was readily distinguishable in the 2D images, enabling relative locations of bound Fabs to be estimated (Fig. 2E). Notably, in nearly all 2D class averages of protomers there was a Fab bound to the base (Fig. 2,B to F).

In some studies, we observed almost no intact trimers with overnight incubation, although shorter incubation times with Fabs (30-60 minutes) could be used to increase the percentage of intact trimers. When rh2688 serum Fabs were added to the MT145KdV5 SOSIP.664 Env trimer(*27*) for 30 minutes and then deposited on an EM grid, base binding antibodies bound to trimers were visible in 2D classes, but protomers were not yet observed (Fig. 2B). For comparison, the same sample when incubated overnight, resulted in extensive trimer disassembly (Fig. 2C). Notably, trimers alone when incubated overnight do not decompose into protomers (Fig. 2A). These observations are consistent with a mechanism in which antibody binding drives the degradation process.

Close inspection of some 2D images revealed that Fabs were bound to both sides of the Env protomer (Fig. 2C, D, F), suggesting that at least some Fabs recognized epitopes not exposed on an intact Env trimer. For B cells to gain access to those off-target neo-epitopes and generate an antibody response, the trimer must therefore degrade in vivo. While the 2D images are quite revealing we could not generate stable 3D reconstructions of the protomers, consistent with a high degree of heterogeneity.

### Antibody-dependent trimer degradation increases over the course of vaccination

In our previous EMPEM study of serial vaccinations in rabbits, we showed a progression in antibody responses starting with base-binding antibodies, with some animals going on to develop neutralizing antibodies to the glycan hole epitope(*25*). Based on the observations described above, we went back and reanalyzed these data to investigate whether trimer degradation also occurred in that experiment. Because we had collected data after the priming and each of the boosts, we were also able to monitor how trimer degradation evolved over time. Two weeks after the priming vaccination using BG505 SOSIP.664 Env, nsEM showed either unliganded Env or base-binding antibodies bound to trimer (Fig. 3A and fig. S7). At 6 weeks post immunization and 2 weeks after a boost, nsEM showed base-binding antibodies and protomers with Fabs bound (Fig. 3B and fig. S8). A cartoon depiction (Fig. 3E) shows the progress of trimer disassembly in vivo. Base-binding antibodies appear first but may not necessarily cause the trimer to disassemble. As the immune response progresses, base-binding antibodies bind at higher stoichiometry and drive the trimer disassembly phenotype. The resulting protomers then present internal epitopes to the immune system and elicit additional off-target responses. In some cases, the secondary antibody responses are quite extensive and 5-6 Fabs can be visualized bound to a single protomer (Fig. 2,C to D, 3C to D and fig. S9). These undesirable epitopes appear highly immunogenic and may therefore distract the immune system from focusing on the more desirable protective broadly neutralizing epitopes.

**Figure 3.**
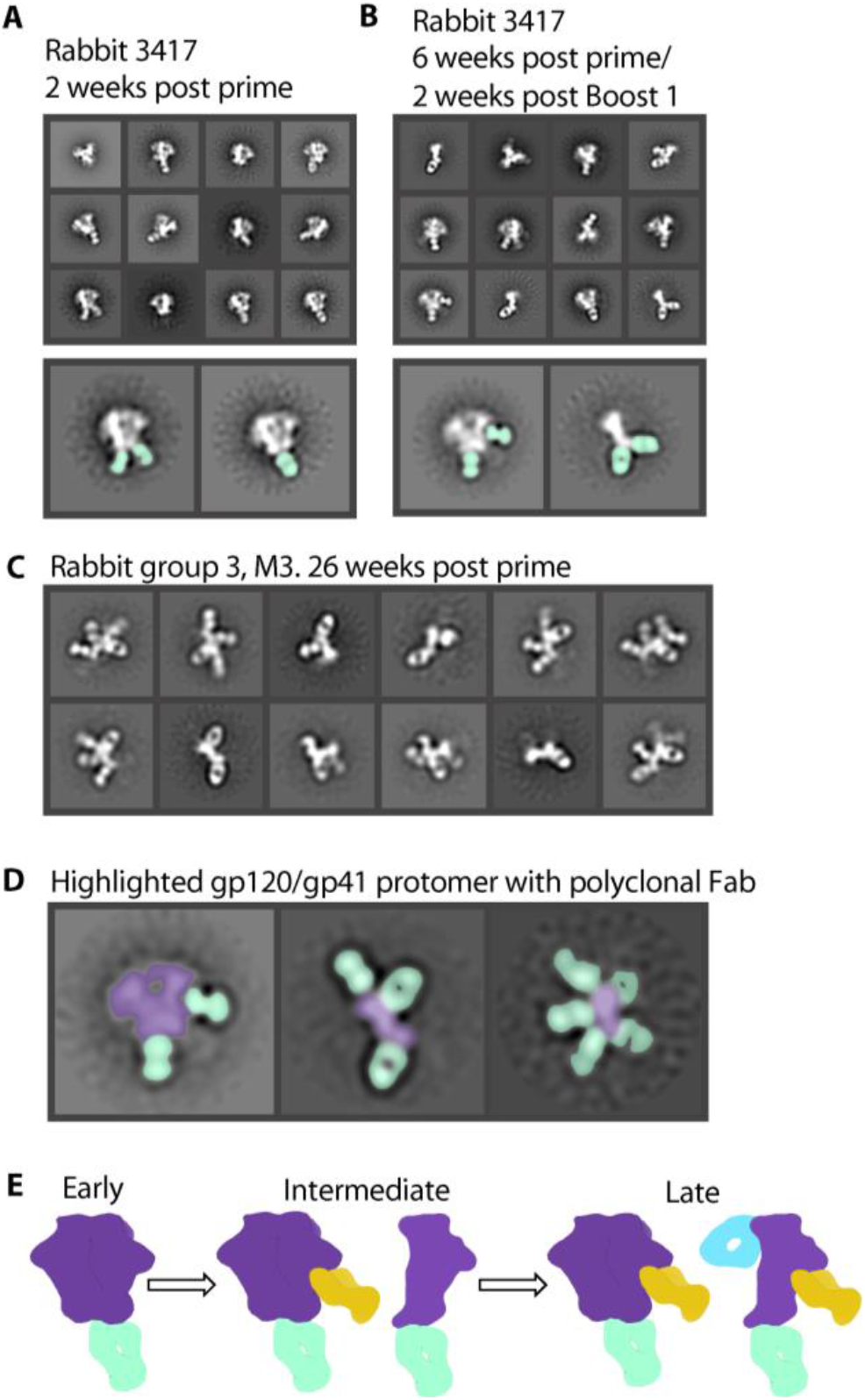
Antibody-dependent trimer degradation occurs over time. (**A**) Two weeks after initial injection of antigen, the immune system develops base binding antibodies but Env stays intact. (**B**) 6 weeks post prime and 2 weeks after a boost, the immune system develops new glycan hole peptide region antibodies and base binding antibodies which cause trimer degradation. (**C**) 26 weeks after initial injection, rabbit serum causes complete degradation of trimer with Fabs bound to all sides of protomer. (**D**) Highlighted classes show intact Env trimer (purple) with Fabs (seafoam green) and protomers (purple) decorated with Fabs. (**E**) Early timepoints show immediate base response (seafoam green). As time goes on, other potentially neutralizing antibodies appear (mustard yellow). At the same time, the base response causes degradation of the Env trimer, resulting in off-target secondary responses (blue).

### Broadly neutralizing monoclonal antibodies can also cause trimer degradation

In previous studies, we also observed timer degradation into protomers in the presence of the bnAbs 3BC315 and 1C2(*4*, *29*). Both antibodies bind the base of the Env trimer and interact with gp120 and gp41 (Fig. 4). A long HCDR3 wedges itself into the gp120/gp41 interface and disrupts the tryptophan clasp region. This insertion causes a conformational change in gp41 that leads to destabilization and degradation. RM20C, which was recently isolated from a BG505 SOSIP.664-immunized macaque, binds to the base of the trimer and causes degradation into protomers when incubated overnight (Fig. 5A and fig. S10). Interestingly, RM20C does not neutralize BG505 virus as an IgG but can weakly neutralize as a Fab, consistent with the hypothesis that the disposition of the second Fab arm and Fc can sterically inhibit access to the base epitope on intact virions(*26*). Therefore, although the base epitope shared by two known bnAbs and one autologous neutralizing Fab, is a bona fide site of vulnerability, access is highly constrained. Binding to this epitope in the context of the soluble trimer immunogen does not have the same constraints. Thus, non-neutralizing antibodies that approach the trimer at a steeper angle than 3BC315 and 1C2, which approach parallel to the membrane, or have the heavy chain of the antibody positioned lower than the light chain, which is likely the case for RM20C, are most likely the driving force in polyclonal serum that causes trimer degradation(*26*).

**Figure 4.**
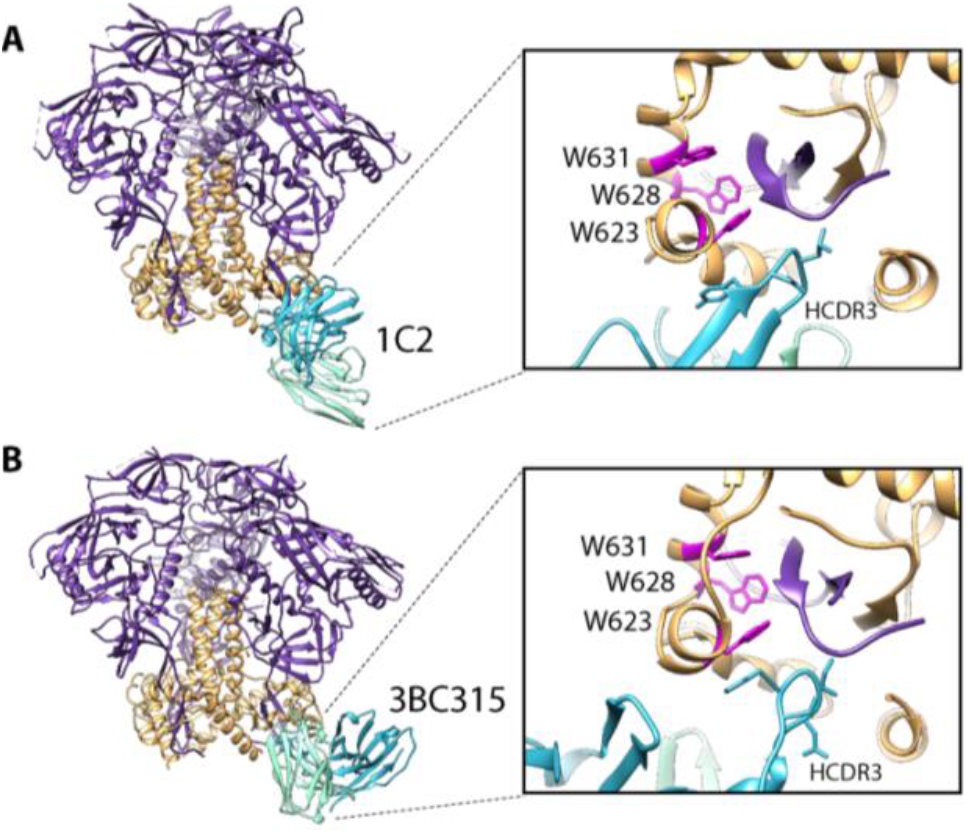
Trimer degradation caused by monoclonal antibodies. (**A**) Structure of 1C2 antibody bound to 16055 NFL TD 2CC +. Env gp120 in purple, gp41 in orange. (**B**) CryoEM derived structure of BG505 SOSIP.664 in complex with 3BC315. Zooming in on the epitope/paratope shows long HCDR3 (blue) wedge up into the base of the trimer and disrupt the tryptophan clasp (magenta) causing degradation.

**Figure 5.**
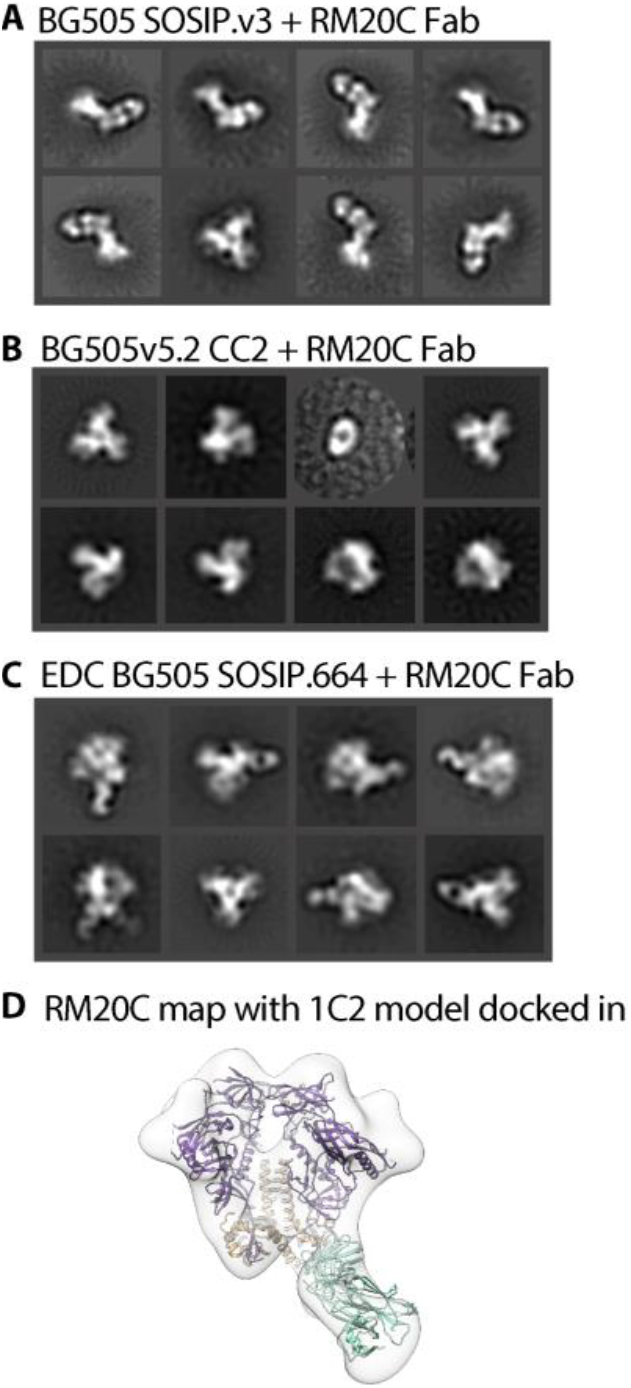
Trimer degradation causing Fab in complex with BG505 variants. (**A**) nsEM 2D classes shows that BG505 SOSIP.v3 degrades into protomers when incubated overnight with RM20C. (**B**) Adding a CC mutation prevents binding of the RM20C Fab all together. (**C**) Chemically cross-linking EDC BG505 SOSIP.664 keeps trimer intact when RM20C binds. (**D**) nsEM derived map of RM20C in complex with EDC BG505 SOSIP.664. Model of 1C2 (6P65, purple, orange) docked in with 6PEH (seafoam green) docked in as Fab.

### Additional disulfide bonds and chemical crosslinking prevent trimer degradation

While antibody-induced trimer degradation appears to be a problem for many of the current Env trimers being used for immunogenicity studies, there are versions of the trimer resistant to this phenotype. We recently demonstrated that an advanced version of the ConM SOSIP trimer (ConM SOSIP.664.v9) that includes an engineered inter-protomer disulfide bond between residues 72 and 564 resists serum-induced degradation that was observed in the parental ConM SOSIP.v7(*21*). An inter-protomer disulfide between residues 501 and 663 of the 16055 Env was also recently reported to prevent native flexibility linked (NFL) trimer degradation induced by bnAbs 3BC315 or 1C2(*4*, *30*). This disulfide bond is an alternative to the traditional 501 to 605 disulfide present in previously described SOSIP trimers. Here we introduced this disulfide bond into the fully-cleaved version of the BG505 Env trimer (referred to as BG505v5.2 CC2). The BG505v5.2 CC2 trimer binds to 3BC315, but resists trimer degradation following overnight incubation. Additionally, BG505-CC2 could bind sCD4, but did not undergo a CD4-induced conformational change(*4*). The CC2 disulfide also knocked out binding to RM20C Fab (Fig. 5B and fig. S11).

Chemical cross-linking, is another potential remedy for trimer disassembly. To test this we incubated BG505 SOSIP.664 trimers that had been crosslinked using 1-ethyl-3-(3-dimethylaminopropyl)carbodiimide (EDC)(*31*) with serum from BG505 SOSIP.664 immunized macaques. The parental BG505 SOSIP.664 trimer readily degrades into protomers while the EDC crosslinked trimers remained resistant to disassembly. EDC crosslinked trimers could also bind RM20C Fab, but resisted degradation following overnight incubation (Fig. 5C, D and fig. S12, S13).

## Discussion

On the surface of the virion, Env is membrane bound. Despite close juxtaposition to the membrane epitopes near the base of the trimer (e.g. 3BC315, 35022, and 1C2)(*4*, *22*, *23*) and the highly conserved membrane proximal external region (MPER)(*32*, *33*), are accessible to neutralizing antibodies. Soluble Env trimers truncated at residue 664 expose a large peptidic surface at the base of the trimer that is not accessible on membrane bound Env. Without the membrane or gp41 glycans, the exposed base has emerged as an easy target for antibodies and therefore of the immunodominant response in different animal models(*6*, *7*, *18*, *19*).

Given the high thermal stability of these trimers and elicitation of neutralizing antibodies, it has been assumed that the trimers stay intact in vivo. Here, we show that, in addition to being off-target and immunodominant, base-directed responses that bind at or around the Trp clasp induce disassembly of the trimer into protomers. Somewhat ironically this immunodominant response targets what can be a bnAb epitope, but one which is highly sensitive to the angle of approach and/or heavy/light chain disposition. Dissociated protomers then become immunogenic in their own right, eliciting a second wave of non-neutralizing antibodies that target epitopes that reside inside the native trimer. As these epitopes are relatively large continuous peptidic surfaces devoid of glycans, like the base of the trimer, they are easier targets for recognition by B-cells than the highly glycosylated exterior surface of the trimer. Thus, the immunogenicity of the trimer base represents a doubly confounding effect on neutralizing antibody responses to the trimer. While trimer degradation has occurred in all trimers that we have analyzed so far, the susceptibility and timing of degradation varied by subtype and design. Some, like BG505 SOSIP.664, are less susceptible while others, such as ConM SOSIP.v7, MT145KdV5 or CRF250 SOSIP.664(*27*, *34*), are much more prone to antibody-dependent degradation. Antibody-induced degradation of trimers may also have implications for binding and antigenicity studies such as Octet or ELISA and may therefore confound interpretation.

Encouragingly, Env trimer constructs already exist that contain inter-protomer disulfide bonds that are resistant to antibody-dependent degradation(*21*). Additional disulfides may provide further stability, although these will need to be tested in immunogenicity studies. Thus, secondary immune responses to internal protomer epitopes may not be an issue provided more advanced designs are utilized. Currently, all of the Env trimers that are most advanced in preclinical or clinical programs (ClinicalTrials.gov: NCT03699241, NCT04177355, NCT03783130) do not have such stabilizing disulfides and we predict that humans will therefore also produce base-directed trimer degrading antibodies. Chemically cross-linked trimers, which we demonstrated are resistant to disassembly, are also being tested in humans by the European AIDS Vaccine Initiative (eavi2020.org).

Attempts to mask the base of the trimer via attachment to nanoparticles or alum have only had marginal impact on reducing base-directed antibodies(*20*, *21*). Thus, further development of these strategies will be required, likely to include inter-protomer disulfide bonds. There is also the potential for prime boost immunization strategies using Env trimers of different sequences at the base such that they cannot be recognized (and degraded) by the base-directed antibodies elicited by the prior immunization. Because the base of the trimer is typically shielded by the membrane it has a relatively high degree of sequence conservation, which makes this strategy less viable. Hence, further mutating exposed base residues, or resurfacing, may be required for such an approach to work. Display of Env trimers on liposomes or delivered via nucleic acid and expressed on cell surfaces in vivo may also limit the base-directed responses.

In summary, our study reveals a previously unappreciated phenomenon of base-directed antibody-induced in vivo Env subunit trimer degradation and secondary antibody responses. These phenomena are unlikely to be unique to HIV Env. Given large number of efforts to employ subunit vaccines against other pathogens including influenza, Respiratory Syncitial Virus (RSV)(*35*), SARS-CoV-2, and others, it will also be important to assess antibody mediated disassembly in those systems as well. EMPEM was integral to our ability to uncover this confounding factor, further demonstrating the power of single particle EM approaches to comprehensively characterize heterogenous ensembles of proteins and protein-protein complexes. Importantly, standard single particle EM workflows usually eliminate, or overlook, data that does not contribute to stable 3D reconstructions. Hence, we advise that future studies include comprehensive 2D and 3D classification to reveal the full extent of species in a given ensemble of Env-antibody complexes.

## Materials and Methods

### Ethics Statement

The rabbit and rhesus macaque animal studies were approved and carried out in accordance with protocols provided to the Institutional Animal Care and Use Committee (IACUC) respectively at The Scripps Research Institute (TSRI; La Jolla, CA) under approval number 19-0020 and at Wisconsin Primate Center (WPC) under approval number G005109. The animals at both facilities were kept, immunized, and bled in compliance with the Animal Welfare Act and other federal statutes and regulations relating to animals and in adherence to the Guide for the Care and Use of Laboratory Animals (National Research Council, 1996).

### Immunization and Sampling

24 female 12-week-old New Zealand white rabbits were prime (wk0: CRF250 SOSIP trimer) and boosted (boost-1: wk8 (CRF250 SOSIP (group-K), or BG505-CRF250V1V2 chimeric SOSIP (group-L) or its N130 (group-M) or V2’ (group-N) glycan filled SOSIP trimers) and boost-2: wk24 (ZM179-ZM233V1V2 chimeric SOSIP trimer) with 50ug of trimer protein and 375 μg of ISCOMs-like saponin adjuvant in 600 uL PBS (300 uL IM injection into each hind leg) (Song et. al. in preparation). Blood was drawn from the marginal ear vein into EDTA or untreated blood collection tubes from select time points (wk2, wk8, wk10, wk26) and the polyclonal IgGs for EMPEM were purified from immune sera/plasma. All procedures were performed in anesthetized animals. 2 groups of 6 rhesus macaques (RMs) each, all females between the ages 5-7 years, were immunized with chimpanzee SIV MT145KdV5 SOSIP (group-1) or MT145KdV5 SOSIP conjugated with alum through pSer peptide linkers (Song et. al. in preparation), as described previously(*20*). Animals were prime (wk0) boosted (wk8) intramuscularly by injecting 100ug of MT145KdV5 or MT145KdV5-pSer trimer proteins along with 1mg alum and 375ug of SMNP adjuvant per animal per immunization. EDTA blood at select time points was collected from animals for isolating polyclonal IgGs for EMPEM analysis.

### Monoclonal Fab Production

RM20C Fab was expressed in HEK293F cells and purified using affinity chromatography. Briefly, HEK293F cells (Invitrogen) were co-transfected with heavy and light chain plasmids (1:1 ratio) using PEImax. The transfection was performed according to the manufacturer’s protocol. Supernatant was harvested 6 days following transfection and passed through a 0.45 μm filter prior to purification using CaptureSelect™ CH1-XL (ThermoFisher) affinity chromatography.

### Env Protein Production

To produce the BG505-CC2 construct, the following mutations were introduced into the BG505 SOSIPv5.2 construct(*17*) using the QuikChange Lightning Multi Site-Directed Mutagenesis Kit (Agilent): P240T, S241N, M271I, F288L, T290E, P291S, R500A, L568D, V570D, R585H, C605T, S613T, Q658T and L663C. Env trimers were expressed in 293F cells and purified using PGT145 affinity chromatography as described previously(*17*).

### Env controls and monoclonal complexes

Unliganded Env trimers were deposited directly onto nsEM grids as controls. Env trimers were incubated overnight at room temperature with monoclonal Fabs in excess of 4.5 molar. Complexes were diluted to 0.03mg/mL and deposited onto nsEM grids. EDC BG505 SOSIP.664 trimer was purified and chemically crosslinked as described previously (*29*, *31*).

### EMPEM

The polyclonal IgGs from serum or plasma were purified by incubating with equal volume of Protein A and G sepharose resin (GE Healthcare) at 4°C overnight, followed by washing with PBS and elution of IgGs with 2M Citric Acid into 0.2M TrisBase. Eluents were buffer-exchanged into PBS using 30kD Amicon tube protein concentrators (MilliporeSigma). The purified polyclonal IgGs were then digested into Fabs using Pierce Fab Preparation Kit (Thermo Scientific). Briefly, after preparation of the IgG sample using Zeba Spin Desalting Columns, 0.25ml equilibrated Immobilized Papain were incubated with 1-2 mg of IgG in digestion buffer rotating for 4-5 hours at 37°C. Digest was separated from Immobilized Papain by centrifugation, followed by mixing incubation with Protein A resin at room temperature for 30min. The flow-through after centrifugation containing Fab fragments were then buffer-exchanged into TBS using 10kD filter Amicon tubes (MilliporeSigma). For EMPEM, 500ug of serum Fab was added to 15ug of Env trimer and incubated overnight at 4C. The EMPEM complex was purified from excess Fab using size exclusion chromatography. The complex peak was concentrated to about 0.03mg/mL and deposited directly onto a nsEM grid.

### Negative stain electron microscopy

Env complexes were added to carbon-covered 400 mesh copper grids. Grids were stained with 2% UF. Micrographs were collected using Leginon(*36*) on either a TF20 or Spirit electron microscope. They were equipped with either a TVIPs or EAGLE 4kx4k camera. Particles were picked using DoGpicker and stacked using Appion(*37*). Particles were 2D classified by Relion (2 and 3) (*38*, *39*). Particles resembling an Env trimer or Env protomer were selected for further processing. Coloring of 2D class images was done in Photoshop. 3D maps were visualized in Chimera(*40*).

## Acknowledgements

We are grateful to Bill Anderson for expert microscopy assistance, JC Ducom and Charles Bowman for computational support, Lauren Holden for editing, and Gabriel Ozorowski for data processing support.

## Author Contributions

Conceptualization (HLT, ABW), Methodology (HLT, ABW), Validation (RA), Formal analysis (HLT, ABW), Investigation (HLT, RA, GS, SC, FA, TJM, MM, MS, EGR), Data curation (HLT), Writing - original draft (HLT, ABW), Writing - review & editing (RA, DRB, CAC, QJS), Supervision (DRB, ABW), Project administration (ABW), Funding acquisition (DRB, ABW, RA). All authors were asked to comment on the manuscript. MS prepared ISCOMs-like saponin adjuvant. WA and MM prepared pSer-linked MTK145 trimers. QJS provided EDC crosslinked trimers.

## Data and materials availability

All data needed to evaluate the conclusions in the paper are present in the paper and/or the Supplementary Materials. Negative stain electron microscopy reconstruction has been deposited in the Electron Microscopy Databank (accession number: EMD-23373).

