## Supplementary Materials for "Disassembly of HIV envelope glycoprotein trimer immunogens is driven by antibodies elicited via immunization"

Published:

The PDF file includes:

Figs. S1 to S13.

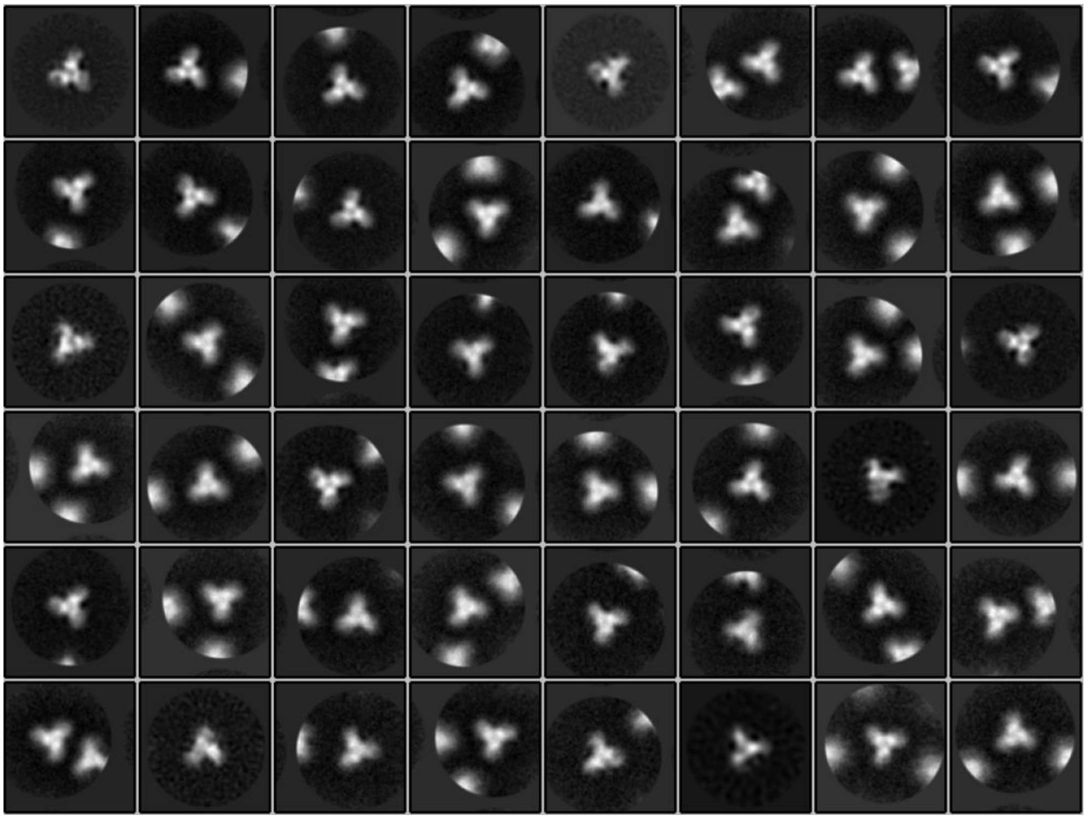

**Fig. S1. 2D classes of unliganded MT145KdV5 trimer.** Only top views due to orientation bias without ligand.

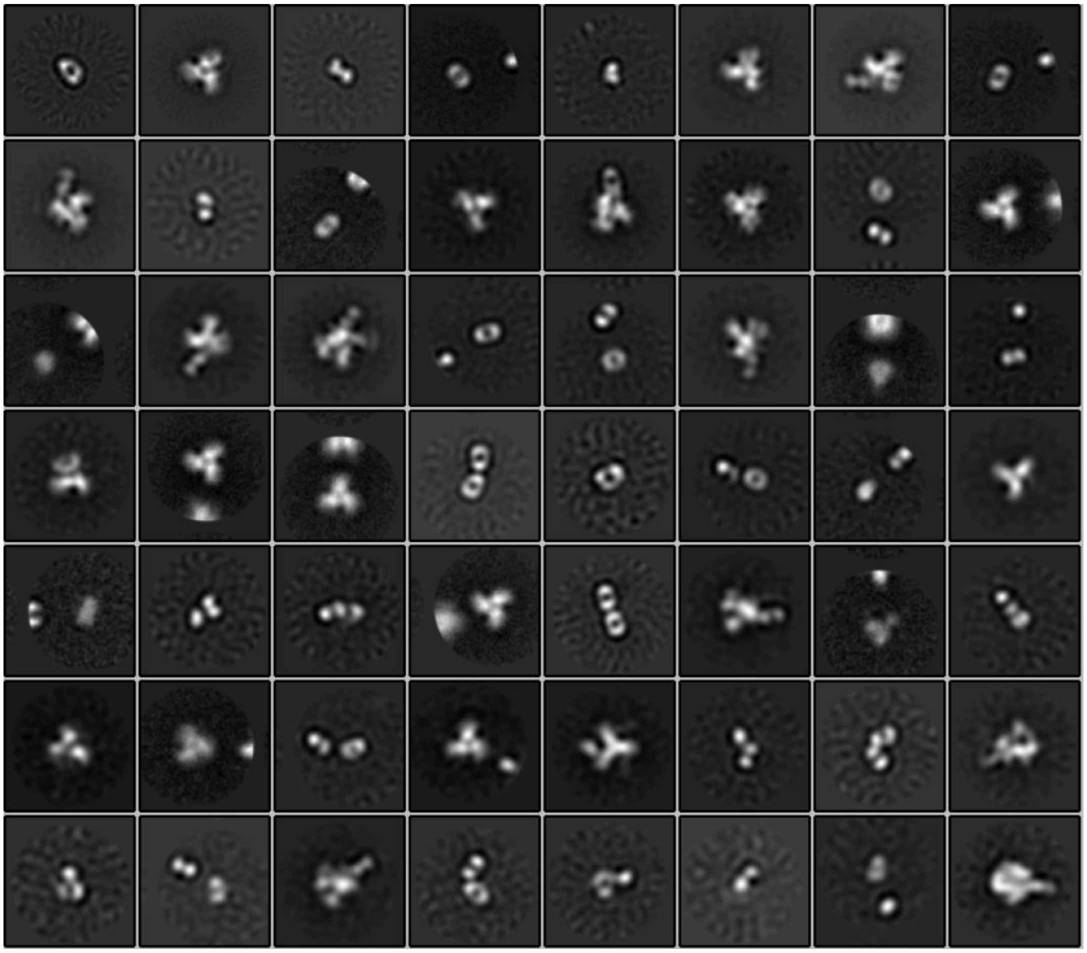

**Fig. S2. 2D classes of rhesus macaque 2688 serum in complex with MT145KdV5, incubated for 30 minutes.** Initial 2D classes of rhesus macaque 2688 fab-isolated serum 10 weeks after immunization with MT145KdV5 SOSIP pSer. Fab in complex with MT145KdV5 SOSIP.664 and incubated for 30 minutes before being added to nsEM grid.

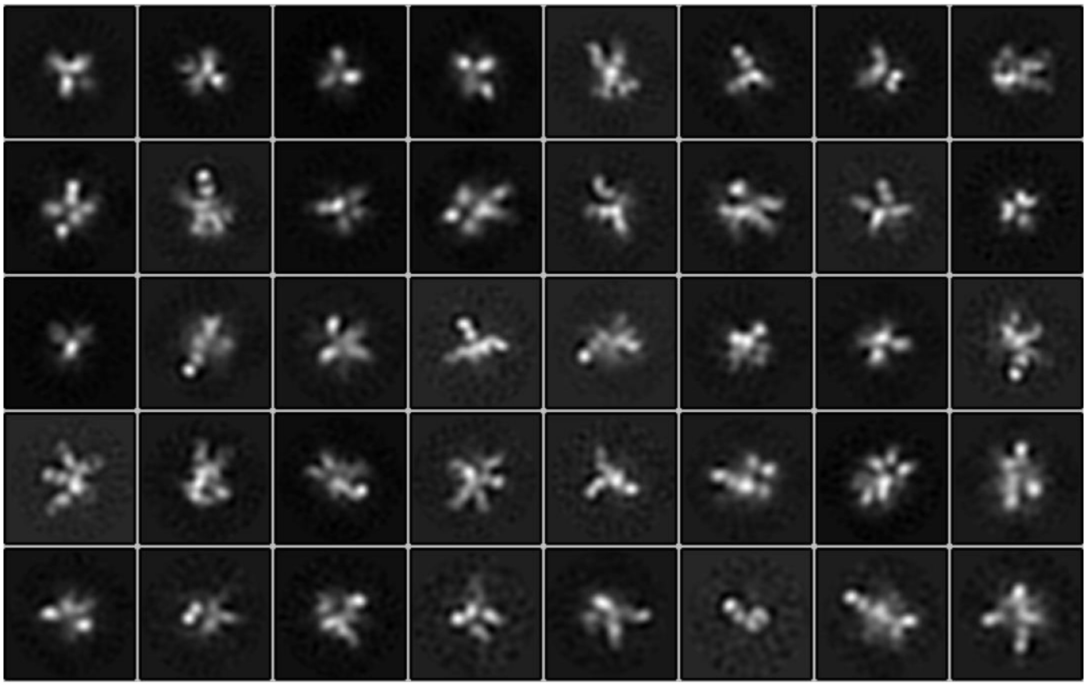

**Fig. S3. 2D classes of rhesus macaque 2688 serum in complex with MT145KdV5**. Initial 2D classes of rhesus macaque 2688 fab-isolated serum 10 weeks after immunization with MT145KdV5 SOSIP pSer. Fab serum in complex with MT145KdV5 SOSIP.664 and incubated overnight.

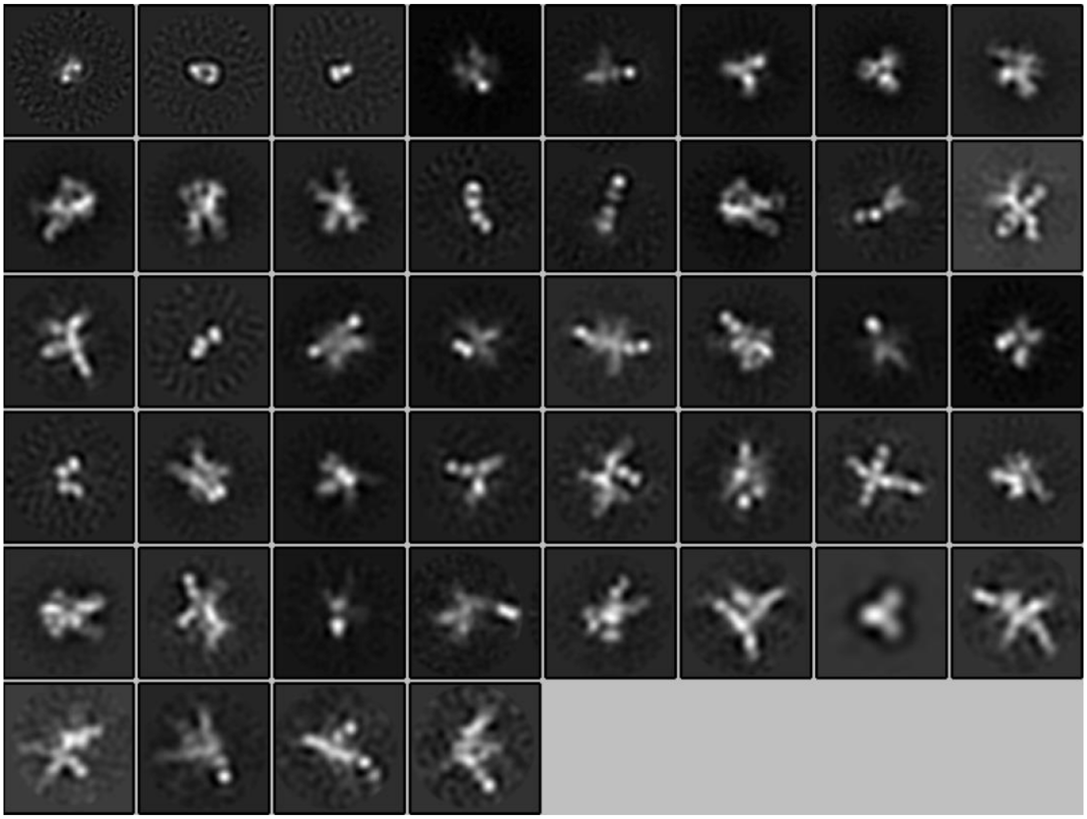

**Fig. S4. 2D classes of rhesus macaque BK89 serum in complex with MT145KdV5 SOSIP.664.**
Initial 2D classes of rhesus macaque BK89 fab-isolated serum in complex with MT145KdV5 SOSIP.664, 10 weeks after immunization with MT145KdV5 SOSIP.664.

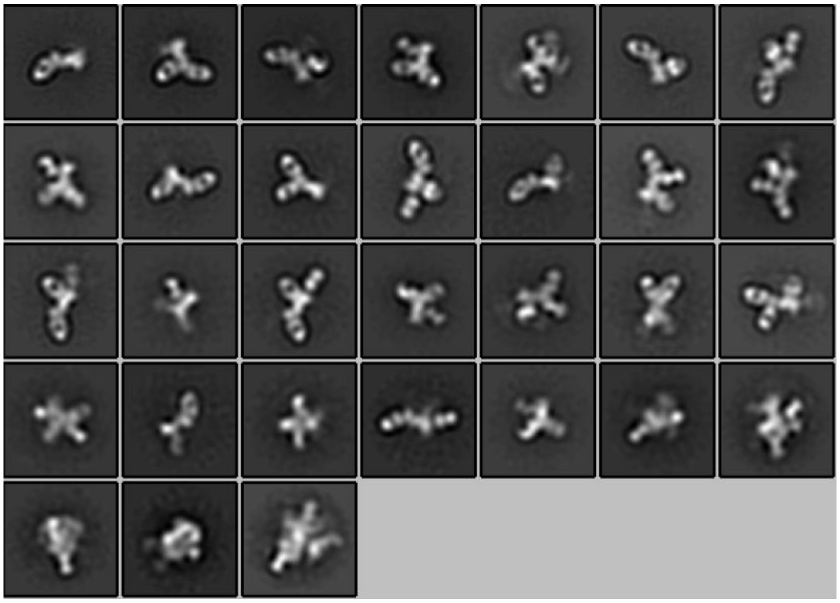

**Fig. S5. 2D classes of rabbit K3 serum post 26 weeks immunization.** Initial 2D classes of rabbit K3 fab isolated serum after 26 weeks of immunization with BG505-CRF250V1V2-grafter trimer. Fab serum in complex with CRF250 SOSIP.664 Env trimer.

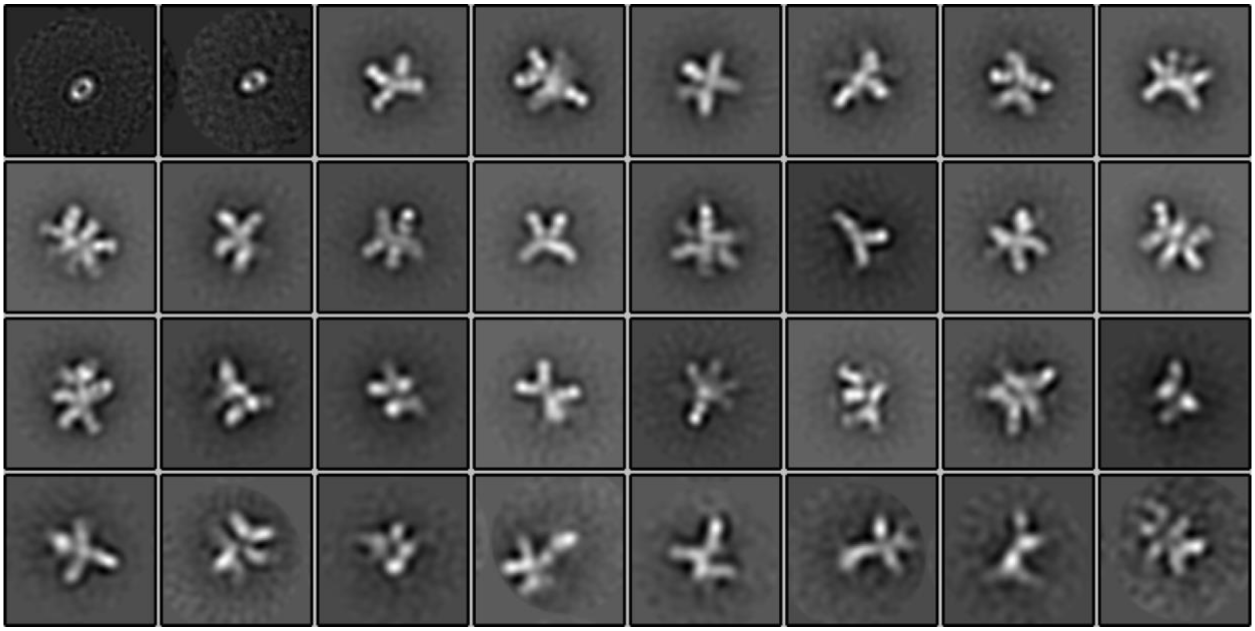

**Fig. S6. 2D classes of rabbit 2382 serum immunized with ConM SOSIP.664.** Initial 2D classes of rabbit 2382 fab-isolated serum in complex with ConM SOSIP.664, 22 weeks post immunization with ConM SOSIP.664.

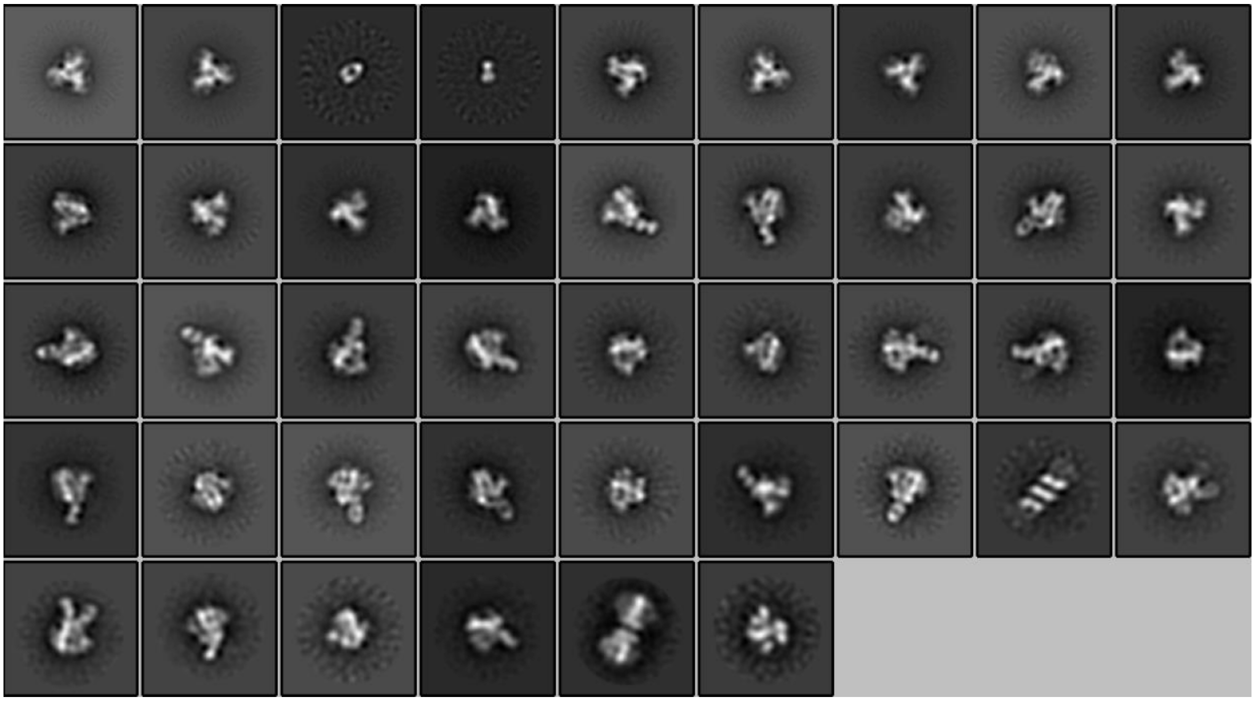

**Fig. S7. 2D classes of rabbit 3417 serum two weeks post prime immunization**. Initial 2D classes of rabbit 3417 fab-isolated serum two weeks after prime immunization of 20ug BG505 SOSIP.664 liposome i.d. Fab serum in complex with BG505 SOSIP.664.

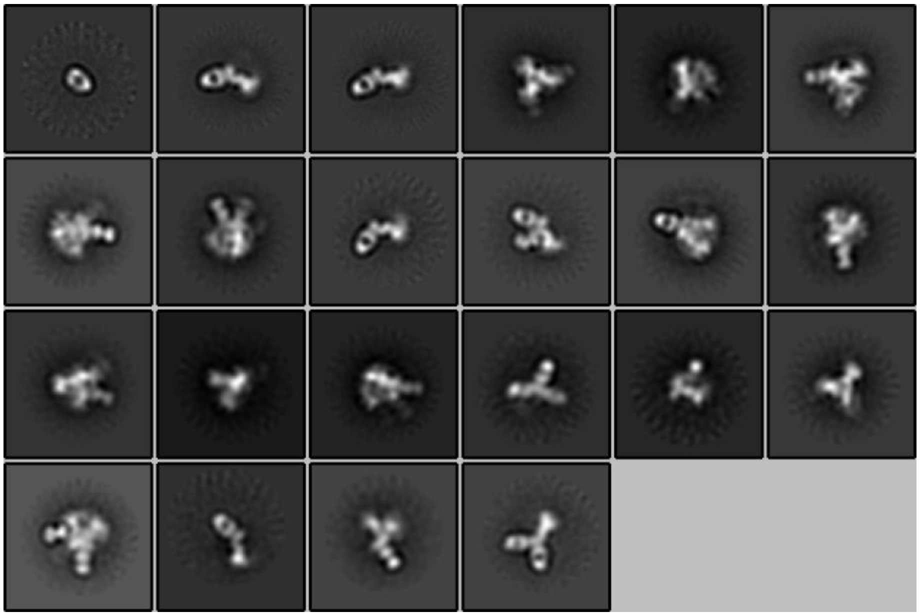

**Fig. S8. 2D classes of Rabbit 3417 after boost 1.** Initial 2D classes of rabbit 3417 fab isolated serum two weeks after 20ug boost 1 via i.m. and 6 weeks after 20ug prime immunization of BG505 SOSIP.664 liposome i.d. Fab serum in complex with BG505 SOSIP.664

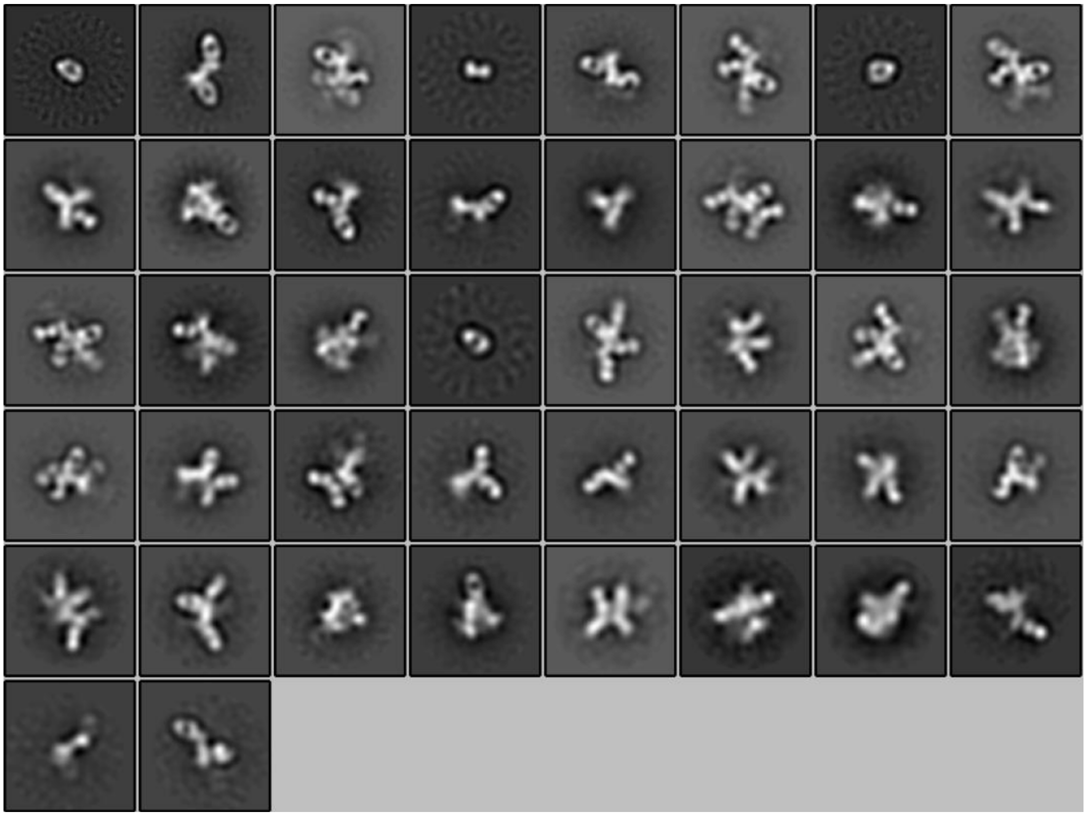

**Fig. S9. 2D classes of rabbit M3 serum 26 weeks after immunization with V1V2-grafted CRF250.** Initial 2D classes of rabbit M3 fab-isolated serum in complex with CRF250, 26 weeks after being immunized with BG505 CRF250V1V2-grafted trimer.

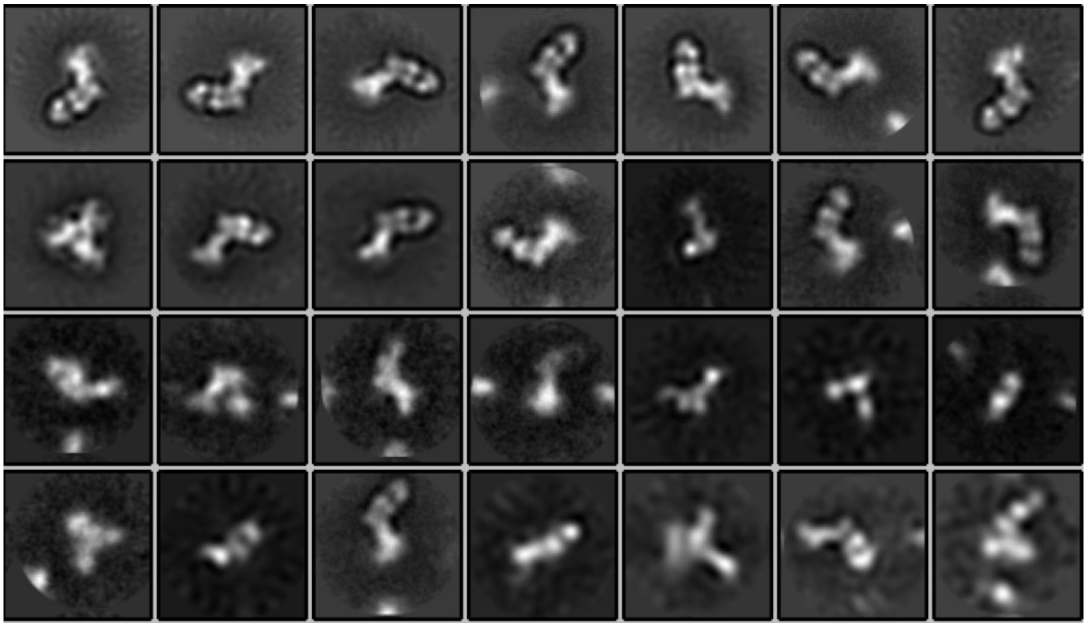

**Fig. S10. 2D classes of RM20C in complex with BG505v3.** Initial 2D classes of RM20C fab in complex with version 3 of BG505 SOSIP.664. Complex was incubated overnight and placed on a nsEM grid.

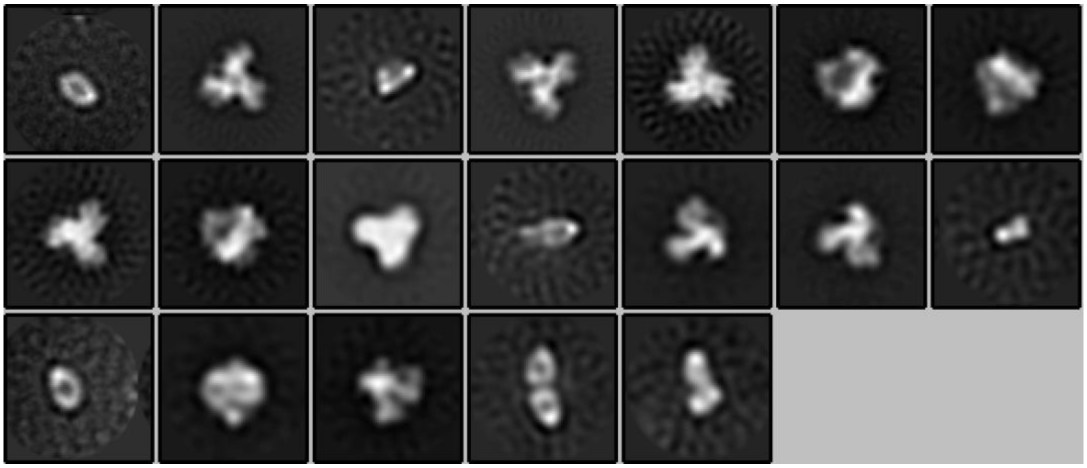

**Fig. S11. 2D classes of RM20C in complex with BG505v5.2 CC2.** Initial 2D classes of RM20C fab in complex with BG505 SOSIP.664 CC2. Complex was incubated overnight and placed on a nsEM grid.

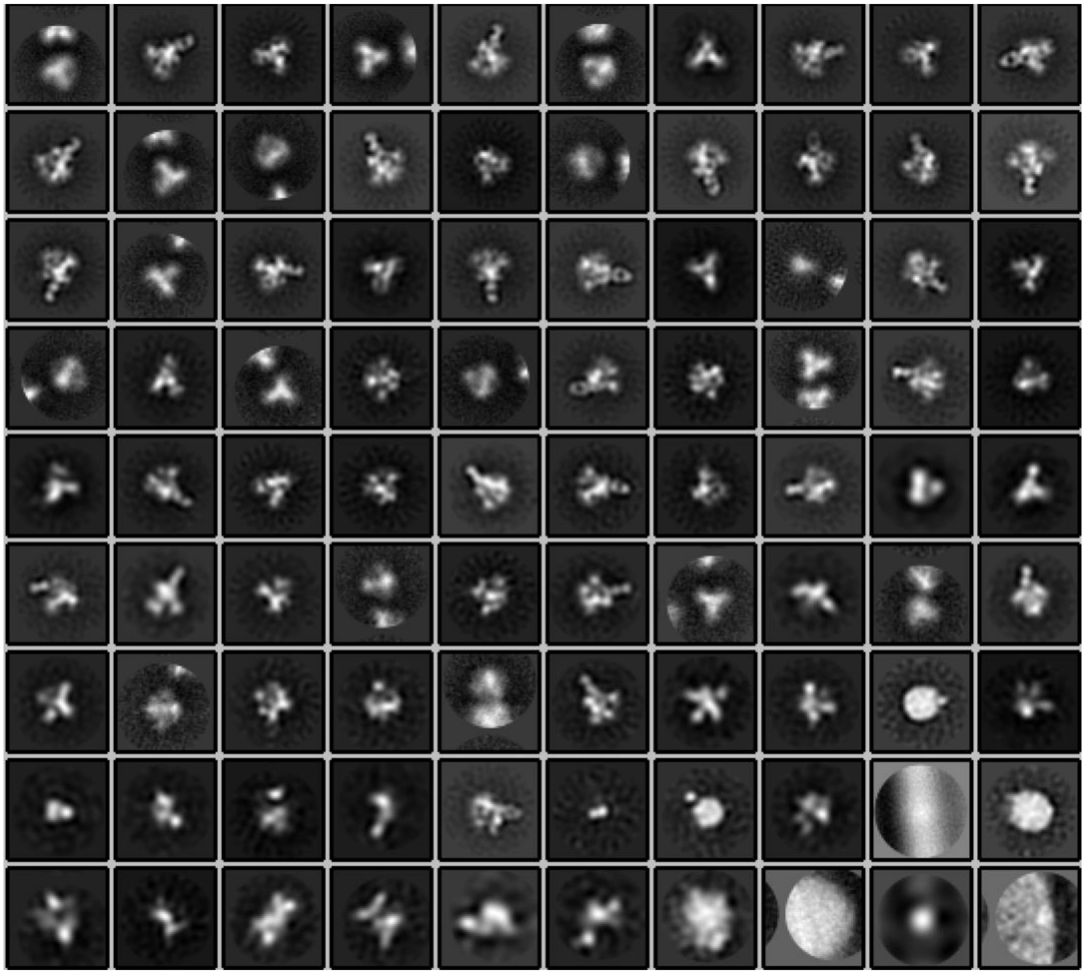

**Fig. S12. 2D classes of RM20C in complex with EDC BG505 SOSIP.664.** Initial 2D classes of RM20C fab in complex with chemical crosslinked (edc) BG505 SOSIP.664. Complex was incubated overnight and placed on a nsEM grid.

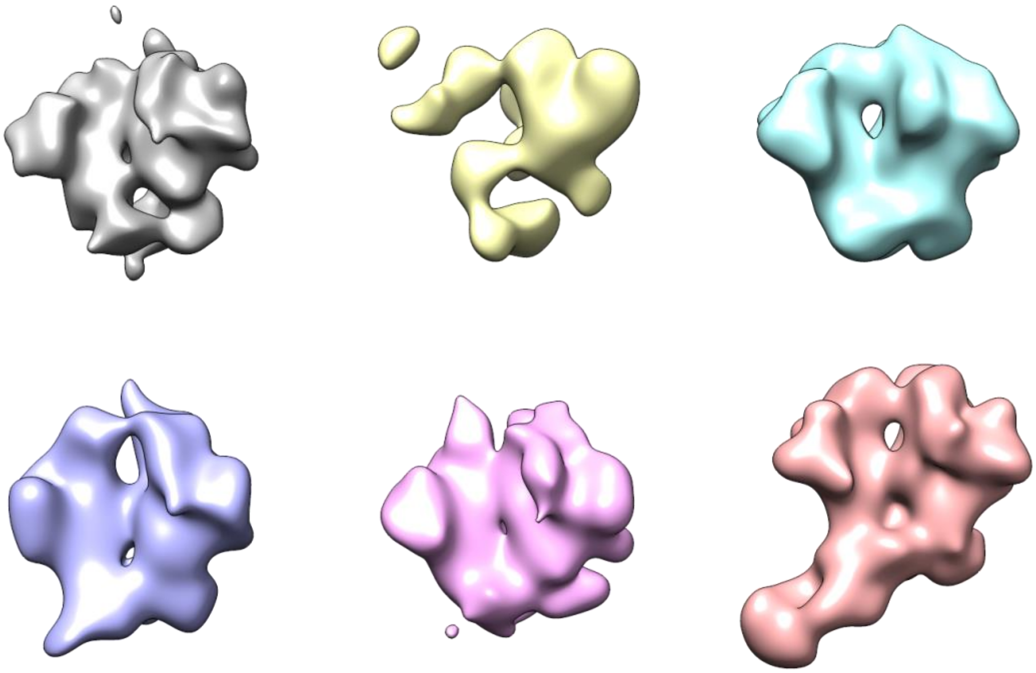

**Fig. S13. 3D classes of RM20C in complex with EDC BG505 SOSIP.664.** Particles from Class 6 (salmon colored) were selected for 3D refinement.
